# First Draft Genome Sequence of the Pathogenic Fungus *Lomentospora prolificans* (formerly *Scedosporium prolificans*)

**DOI:** 10.1101/171975

**Authors:** Ruibang Luo, Aleksey Zimin, Rachael Workman, Yunfan Fan, Geo Pertea, Nina Grossman, Maggie P. Wear, Bei Jia, Heather Miller, Arturo Casadevall, Winston Timp, Sean X. Zhang, Steven L. Salzberg

## Abstract

Here we describe the sequencing and assembly of the pathogenic fungus *Lomentospora prolificans* using a combination of short, highly accurate Illumina reads and additional coverage in very long Oxford Nanopore reads. The resulting assembly is highly contiguous, containing a total of 37,630,066 bp with over 98% of the sequence in just 26 scaffolds. Annotation identified 8,656 protein-coding genes. Pulsed-field gel analysis suggests that this organism contains at least 7 and possibly 11 chromosomes, the two longest of which have sizes corresponding closely to the sizes of the longest scaffolds, at 6.6 and 5.7 Mb.

---

*Lomentospora prolificans* is an opportunistic fungal pathogen that causes a wide variety of infections in immunocompromised and immunocompetent people and animals ^1, 2^. It was originally proposed as *Scedosporium inflatum* in 1984 by Malloch & Salkin ^3^, and later renamed to *Scedosporium prolificans* in 1991 by Geuho and de Hoog ^4^. In 2014, the fungus was re-named *Lomentospora prolificans* based on its phylogenetic distance from other *Scedosporium* species ^5^.

*L. prolificans* is distributed throughout the world and primarily found in soil and plants. Transmission to humans is often via inhalation of the spores produced by the fungus, but occasionally it is introduced by direct traumatic inoculation. Infections can range from mild, local infections to severe and disseminated, with the latter being life threatening. Invasive infections have been increasingly associated with hematological malignancies, transplantation, and cystic fibrosis ^1^. The major challenge to successful therapy is that the fungus is intrinsically resistant to almost all antifungal drugs currently available for treatment ^6^. As a result, the clinical outcome of invasive infections is often fatal.

## Methods

### Isolation and growth

The *L. prolificans* strain included in this study was isolated from a brochoalveolar lavage fluid sample of a cystic fibrosis patient, in whom it was causing a chronic refractory pulmonary fungal infection. The fungus was cultivated on Potato Flake Agar plates (BD, Sparks, MD) at 30°C until reaching sizable colonies with adequate sporulation. Spores were collected and then converted into a hyphal mass by growing in Sabouraud Liquid Broth (BD, Sparks, MD) under a 20-rpm rotator (Stuart, Staffordshire, UK) at 23°C. Genomic DNA was extracted from the hyphal mass by using ZR Fungal/Bacterial DNA MiniPrep kits (Zymo Research, Irvine, CA) according to the manufacturer’s protocol with the following modification: a horizontal vortex adapter (Mo Bio, Carlsbad, CA) was used with the 10-minute beads beating step during the cell lysis step.

### Illumina sequencing library construction

Nextera XT (Illumina) transposase-based libraries were generated with 1 ng of purified, unsheared *L. prolificans* hyphal DNA. After transposition and barcoded adapter ligation by PCR, the library was purified using 0.4X AMPure XP (Beckman Coulter), and size profiles generated using the Agilent Bioanalyzer high sensitivity chips, average size 747 bp. The library was normalized to 4 nM, then paired end, dual index sequencing was performed using Miseq v2 500 cycle chemistry.

### Nanopore sequencing library construction and data preparation

We input 1.5 ug purified, unsheared *L. prolificans* hyphal DNA into the LSK-108 Oxford Nanopore Technologies (ONT) ligation protocol. The library was blunt-ended and A-tailed with NEB Ultra II End prep module, and purified using 1X AMpure XP. Adapter ligation was then performed using Blunt-TA ligase master mix (NEB) and proprietary ONT “1D” adapters containing a pre-loaded motor protein. The library was purified using 0.4X Ampure XP, washed with buffer WB (ONT), and eluted with elution buffer EB (ONT), which contains a tether molecule that directs library molecules towards the nanopore membrane surface. The library was sized using the Agilent Bioanalyzer high sensitivity chips, size peaking at 2.8kb. The entire library was mixed with running buffer and sepharose loading beads (ONT), then run on a R9.4 SpotON MinION flowcell for 48 hours. Raw fast5 files were basecalled using Albacore version 1.0.2, and fastq files were extracted using our custom python script.

### Sequencing

The Illumina MiSeq run generated 12.04 million 250 bp paired-end reads with a mean fragment size of 500 bp, for a total of 6.02 Gbp of data. The Oxford Nanopore MinION run produced approximately 3.66 million reads for a total of 4.3 Gbp of data. Among the MinION reads, 1.18 million reads (2.96 Gbp) were longer than 1 kbp, and 25,788 reads (333.27 Mbp) were longer than 10 kbp.

### Pulsed field gel analysis

Conidia were inoculated into Sabouraud dextrose broth, grown with shaking at 30°C for 2-3 d, and used to generate protoplasts using minor modifications of published methodology ^7^. Briefly, fungal biomass was filtered through a cell strainer, washed with sterile water and incubated at 30°C on a nutating mixer for 3 h in OM buffer with 5% Glucanex. Contents were then split into sterile centrifuge tubes and overlaid with chilled ST buffer in a ratio of 1.2 mL fungal solution to 1 mL ST buffer. Tubes were then centrifuged at 5000 g for 15 minutes at 4°C. Protoplasts were recovered at the interface of the two buffers and transferred to a sterile centrifuge tube, to which an equal volume of chilled STC buffer was added. Protoplasts were pelleted at 3000 g for 10 minutes at 4 °C, following which supernatant was removed, and protoplasts were resuspended in 10 mL STC buffer. This was repeated two more times, with the final re-suspension being performed with 200 μL GMB buffer. Plugs were then made and treated using methodology adapted from Brody and Carbon ^8^. Briefly, 200 μL 1-2 × 10^9^ protoplasts in GMB buffer were mixed with 200 μL 2% low-melt agarose in 50 mM EDTA (pH 8) cooled to 42°C and pipetted into plug molds, which were then placed on ice for 10 minutes to solidify. Plugs were removed from their molds and incubated in NDS buffer with proteinase K at 50°C for 24 hours, followed by three 30-minute washes in 50 mM EDTA (pH 8) at 50°C. Plugs were stored in 50 mM EDTA (pH 8) at 4°C. Plugs were inserted into agarose gels, along with *S. cerevisiae* and *S. pombe* size standards (Bio-Rad Laboratories, Hercules, CA, USA) and clamped homogeneous electrical field (CHEF) electrophoresis was run on a CHEF-DR III (Bio-Rad Laboratories). The gel showing the full range of bands was captured using to the conditions described in the CHEF-DR III manual for *Hansenula wingei* with the following exception: gel was made from 0.8% SeaKem Gold agarose (Lonza, Basel, Switzerland). The gel showing the smaller bands in greater resolution was made using 0.6% SeaKem Gold agarose in 0.5x TBE, with one block of 4.5 V/cm at an included angle of 120° and switch time of 60-300 seconds for 24 hours, followed by a second block of 2.0 V/cm at an included angle of 106° and switch time of 720-900 seconds for 12 hours. The run was conducted at 12°C.

## Results

### Assembly

We used multiple tools in our assembly pipeline (**Figure 1**). Firstly, we used the MaSuRCA genome assembler ^9^ (version 3.2.2_RC3) to assemble both types of data, with default settings except for the option “+USE_LINKING_MATES=1”. Then we used the Megahit genome assembler ^10^ (version 1.1.1) to assemble the Illumina data separately. We then aligned the Megahit assembly to the MaSuRCA assembly using NUCmer (version 3.1) ^11^, and found 2,599 sequences in the Megahit assembly, summing up to 1.01 Mbp, that were missing in the MaSuRCA assembly. We added these sequences to the MaSuRCA assembly to form a more complete set of contigs.

**Figure 1.**
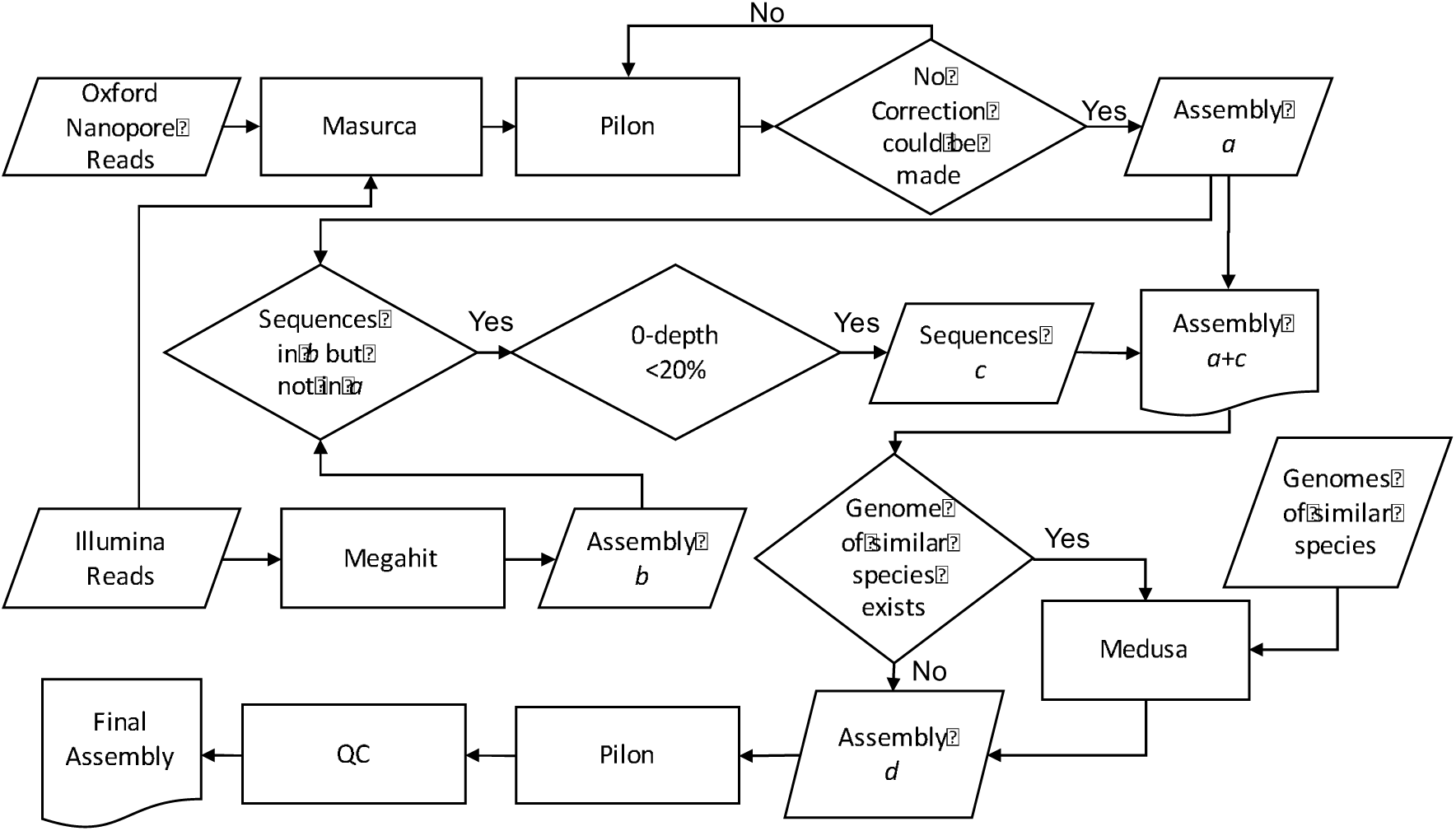
Genome assembly pipeline used for hybrid assembly of *L. prolificans* from Oxford Nanopore and Illumina reads. Both data sets were assembled jointly with MaSuRCA, and Illumina reads were assembled separately with Megahit, followed by assembly polishing, comparison, and merging steps. The genomes of two related *Scedosporium* species, *S. apiospermum* and *S. aurantiasum*, were used to improve scaffolding.

Next we used the MeDuSa scaffolder (version 1.6) ^9^ to determine the correct order and orientation of the contigs using the assemblies of two related organisms *Scedosporium apiospermum* ^12^ and *Scedosporium aurantiasum* ^13^. The resulting scaffolds were polished and gap-filled by Pilon (version 1.5) ^14^ iteratively for 15 rounds until no additional correction could be made. Note that both Illumina reads and MinION reads were used in the first round of polishing, but only Illumina reads were used in the last 14 rounds. Then we aligned the polished scaffolds against the Univec and the Emvec database to ensure no vector sequence was contained in the assembly. Finally, we aligned the Illumina reads and Nanopore reads to the scaffolds using the BWA aligner (version 0.7.15) ^15^. Using these alignments, we broke apart scaffolds at positions with zero coverage, except for the leading and trailing 100 bp. The final assembly consists of 1,629 scaffolds (240 to 6,579,848 bp in length) and has a total size of 37,630,066 bp with 51.46% GC content. The N50 size is 2,796,173 bp. The longest 26 sequences comprise 98.02% of the assembly (Table 1). Four of the longest scaffolds (lengths 6.6 Mb, 1.8 Mb, 876 Kb, and 837 Kb) contain telomeric repeats on one end.

**Table 1.** The size and the cumulative percentage of the total assembly for the 26 longest scaffolds.

| Size (bp) | % | Size (bp) | % | Size (bp) | % |
| --- | --- | --- | --- | --- | --- |
| 6,579,848 | 17.49 | 890,147 | 82.47 | 326,693 | 96.46 |
| 5,696,782 | 32.62 | 878,390 | 84.80 | 173,261 | 96.92 |
| 4,961,516 | 45.81 | 876,043 | 87.13 | 107,743 | 97.21 |
| 2,796,173 | 53.24 | 837,198 | 89.36 | 101,996 | 97.48 |
| 2,583,719 | 60.11 | 594,860 | 90.94 | 89,613 | 97.72 |
| 2,065,651 | 65.60 | 500,465 | 92.27 | 52,380 | 97.86 |
| 1,956,546 | 70.80 | 482,714 | 93.55 | 33,039 | 97.95 |
| 1,805,717 | 75.59 | 431,267 | 94.70 | 26,943 | 98.02 |
| 1,696,794 | 80.10 | 338,834 | 95.60 |  |  |

We used the BUSCO pipeline (v3) ^16^ to assess the genome assembly completeness. BUSCO searches for the presence of genes that occur as single-copy orthologs in at least 90% of a lineage. We used the lineage “Sordariomycetes”, which contains 3,725 orthologous group and is the class containing *Lomentospora prolificans*. Our *Lomentospora prolificans* assembly covers 94.2% of the groups, while the *Scedosporium apiospermum* assembly and the *Scedosporium aurantiasum* assembly cover 94.5% and 90.9% of groups respectively (Figure 2).

**Figure 2.**
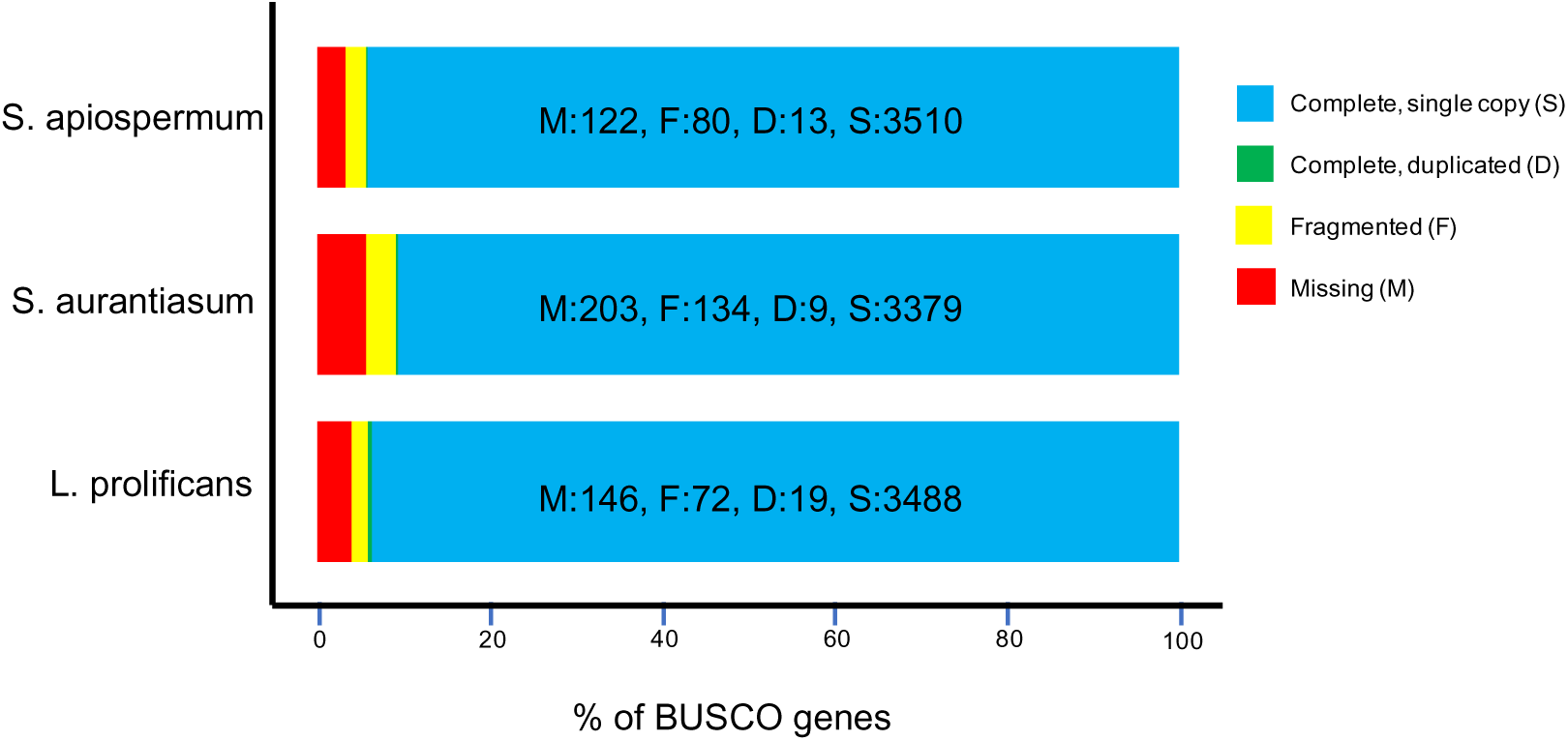
Results from searching for a set of conserved, single-copy genes in L. prolificans (bottom) and its two closest sequenced relatives, *S. apiospermum* and *S. aurantiasum*.

### Annotation

We used the MAKER automated annotation system ^17^ to identify protein-coding genes in the assembly. The primary evidence for annotation was protein and EST alignments from other fungi, which identified 8,539 genes, of which 7,477 were multi-exon transcripts. We then trained three *ab initio* gene finders–Augustus, SNAP, and GeneMarkES–and provided their outputs to MAKER in a second pass. MAKER uses these predictions to modify the alignment-based gene models, although it does not predict any genes based solely on *ab initio* predictions.

After the second pass of annotation, we identified 8,656 genes of which 7,112 contain more than one exon. These transcripts covered 11,657,920 bp, of which 11,643,840 bp are protein-coding and the remainder are 3' and 5' untranslated regions. By comparison, the annotation of the draft genomes of *S. apiospermum* and *S. aurantiasum* contain 10,919 and 10,525 genes respectively. All of these gene predictions are based on automated annotation pipelines and should be regarded as preliminary.

### Chromosome structure

To determine if the assembly was consistent with laboratory estimates of genome size, we separated and visualized the *L. prolificans* chromosomes in agarose by pulsed-field gel electrophoresis (PFGE). While not all chromosomes appeared fully resolved, the lane with the best resolution showed bands for at least 8 chromosomes, at between approximately 1.4 and 6.4 Mb in length (Figure 3A), relative to *S. cerevisiae* and *S. pombe* standards. The chromosome banding pattern above 3.5 Mb was not consistently observed between two sample lanes (data not shown), which may indicate these larger chromosomes were not stable and/or degraded quickly. Of the observed bands, three show staining consistent with multiple chromosomes of the same approximate size migrating together (Figure 3A, bands 1, 4, and 8). It should be noted that the largest two bands were larger than the largest *S. pombe* chromosome, decreasing our ability to predict their sizes.

**Figure 3.**
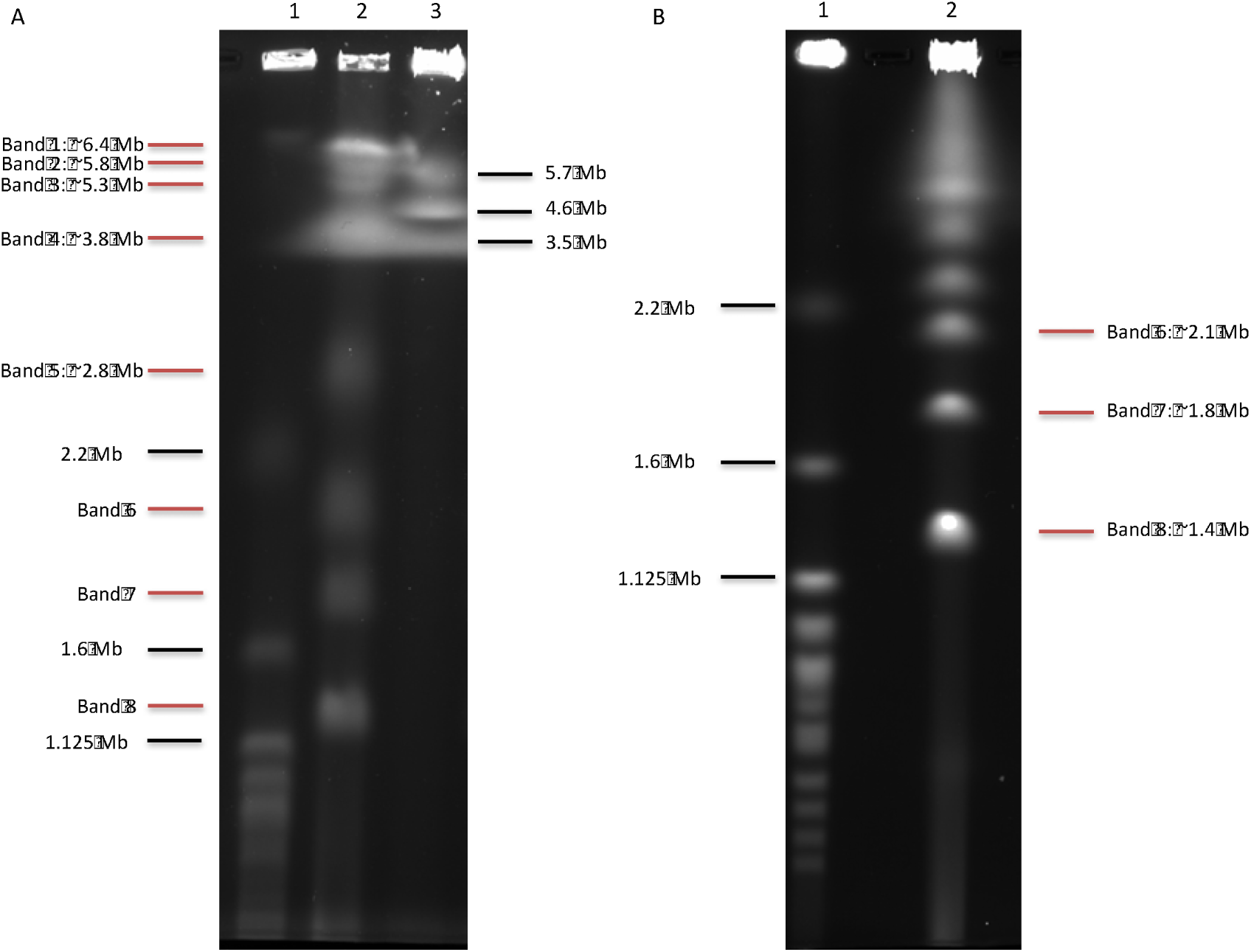
Separation of *L. prolificans* chromosomal DNA by pulsed-field gel electrophoresis. (A) Chromosomal DNA from *S. cerevisiae* (lane 1), *L. prolificans* (lane 2) and *S. pombe* (lane 3) were run, allowing estimation of the number and lengths of the *L. prolificans* chromosomes, as indicated. (B) Chromosomal DNA from *S. cerevisiae* (lane 1) and *L. prolificans* (lane 2) were separated under conditions optimized for chromosomes in the range of 0.9–3.2 Mb, allowing more precise estimates of lengths of chromosomes contained in *L. prolificans* bands 6–8.

A second PFGE (Figure 3B) performed under conditions designed to better resolve bands in the range of 0.9–3.2 Mb allowed us to more precisely estimate the sizes of the three smallest bands, which we estimate to be 1.4, 1.8 and 2.1 Mb, respectively. The 1.4 Mb band (Band 8) showed staining consistent with two chromosomes migrating together. From these gels we can determine that *L. prolificans* contains at least 7, and likely 11 chromosomes. We observe staining consistent with at least two chromosomes at sizes 1.4 Mb, ~3.8 Mb, and ~6.4 Mb. These three sets of two chromosomes, along with those within the range of the standards, 1.8, 2.1, 2.8, 3.8, and 5.3 Mb, plus a single band which is outside of the range, but estimated around 5.8 Mb, sum up to 41 Mb. Given the uncertainty of these estimates, the total approximate genome size is consistent with our 37.6 Mb genome assembly. The two largest scaffolds (Table 1) have lengths 6.6 and 5.7 Mb, which is in agreement with the PFGE results for the largest chromosomes.

### Data availability

This genome project has been deposited at NCBI/GenBank as BioProject PRJNA392827, which includes the raw read data, assembly, and annotation. The assembly is available under accession NLAX00000000; the version described in this paper is version NLAX01000000.

## Acknowledgements

Thanks to Brendan Cormack for help with DNA isolation and preparation, and to Carol Greider and Carla Connelly for assistance with the pulsed-field gel analysis. This work was supported in part by the National Institutes of Health under grants R01-HL129239, R01-HG006677, and T32-AI007417.

